# Efficient semi-supervised semantic segmentation of electron microscopy cancer images with sparse annotations

**DOI:** 10.1101/2023.10.30.563998

**Authors:** Lucas Pagano, Guillaume Thibault, Walid Bousselham, Jessica L. Riesterer, Xubo Song, Joe W. Gray

**Affiliations:** Department of Biomedical Engineering, Oregon Health and Science University, Portland, OR, USA.; Department of Medical Informatics and Clinical Epidemiology at Oregon Health and Science University, Portland, OR USA; Knight Cancer Institute, Oregon Health and Science University, Portland, OR, USA

**Keywords:** Deep learning, semi-supervised, semantic segmentation, electron microscopy, vEM, sparse labels

## Abstract

Electron microscopy (EM) enables imaging at nanometer resolution and can shed light on how cancer evolves to develop resistance to therapy. Acquiring these images has become a routine task; however, analyzing them is now the bottleneck, as manual structure identification is very time-consuming and can take up to several months for a single sample. Deep learning approaches offer a suitable solution to speed up the analysis. In this work, we present a study of several state-of-the-art deep learning models for the task of segmenting nuclei and nucleoli in volumes from tumor biopsies. We compared previous results obtained with the ResUNet architecture to the more recent UNet++, FracTALResNet, SenFormer, and CEECNet models. In addition, we explored the utilization of unlabeled images through semi-supervised learning with Cross Pseudo Supervision. We have trained and evaluated all of the models on sparse manual labels from three fully annotated in-house datasets that we have made available on demand, demonstrating improvements in terms of 3D Dice score. From the analysis of these results, we drew conclusions on the relative gains of using more complex models, semi-supervised learning as well as next steps for the mitigation of the manual segmentation bottleneck.

## 1 INTRODUCTION

Recent advances in cancer nanomedicine have made cancer treatment safer and more effective (Shi et al., 2017). Nanotechnology has elucidated interactions between tumor cells and their microenvironment showing key factors in cancer behavior and responses to treatment (Hirata and Sahai, 2017; Tanaka and Kano, 2018). Gaining a deeper understanding of the underlying mechanisms taking place during such interactions will help us understand how cancer grows and develops drug resistance, and ultimately help us find new, efficient and safe therapeutic strategies aimed at disrupting cancer development (Baghban et al., 2020).

To do this, high resolution information collected from the cellular components at nanometer scale using focused ion beam-scanning electron microscopy (FIB-SEM) is especially useful as it provides volumes of serially-collected 2D SEM images, creating volume EM (vEM) image stacks, and allowing access to 3D information from tissues (Giannuzzi and Stevie, 2005). This fully automated protocol avoids artifacts associated with serial microtomies and enables voxels to be isotropic, thus yielding a similar image quality in all dimensions, beneficial for feature recognition and context within the volume (Bushby et al., 2011).

These advantageous features have made SEM desirable for use in clinical programs. However, the analysis-limiting step is the extraction of meaningful features, starting with the segmentation of cellular components present in these images. This is currently done by human experts through hand annotating. It’s a tedious and time consuming task, making it unsuitable for medical applications and decisions where time is a critical factor. To overcome this limitation and fully make use of FIB-SEM in a clinical setting, the development of automated and robust models is critical to speeding up this task (Perez et al., 2014).

Segmenting images acquired via FIB-SEM is a difficult problem. Indeed, these images differ considerably from natural ones (images representing what human being would observe in the real world), and even from other microscopy techniques such as fluorescence microscopy, due to increased noise, different collection resolution, and the reduced number of image channels. EM images are single-channel (grayscale) and tend to have limited contrast between objects of interest and background (Karabağ et al., 2020). Furthermore, the ultrastructure of tumor cells and their microenvironment vary from those of normal cells (Nunney et al., 2015), and EM analysis methods can be tissue type dependent; most current methods have been developed for neural images (Taha and Hanbury, 2015; Zhang et al., 2021). Therefore, segmentation methods designed to assist other microscopy modalities or other tissue types cannot be applied to ultrastructure segmentation of cancer cells imaged by EM.

We expanded on previous work from (Machireddy et al., 2021), where authors showed that a sparsely manually annotated dataset, typically around 1% of the image stack, was sufficient to train models to segment the whole volume. While state-of-the-art in semantic segmentation has been dominated by attention-based models for natural images (Jain et al., 2021), convolutional architectures remain main stream with EM data, and were used in (Machireddy et al., 2021), and the companion paper within this journal volume. In this paper, we compared architectures as well as training frameworks to find the most suitable one for the task of semantic segmentation in the aforementioned specific context of FIB-SEM images. By optimizing the learning process, we expected to improve overall segmentation results and minimize the manual annotation bottleneck by reducing the number of manually labeled images needed for training.

In this study, we focused on the segmentation of nuclei and nucleoli in vEM image stacks acquired from human tumor samples, as both are commonly used as cancer cell identifiers (Zink et al., 2004) and have emerged as promising therapeutic targets for cancer treatment (Lindström et al., 2018). Segmenting both structures accurately has thus proven essential. We evaluated the selected models on FIB-SEM images of three longitudinal tissue biopsy datasets that are available on demand as part of the Human Tumor Atlas Network (HTAN). A quick visualization of the data and end results can be found in Figures 1 and 2.

**Figure 1.**
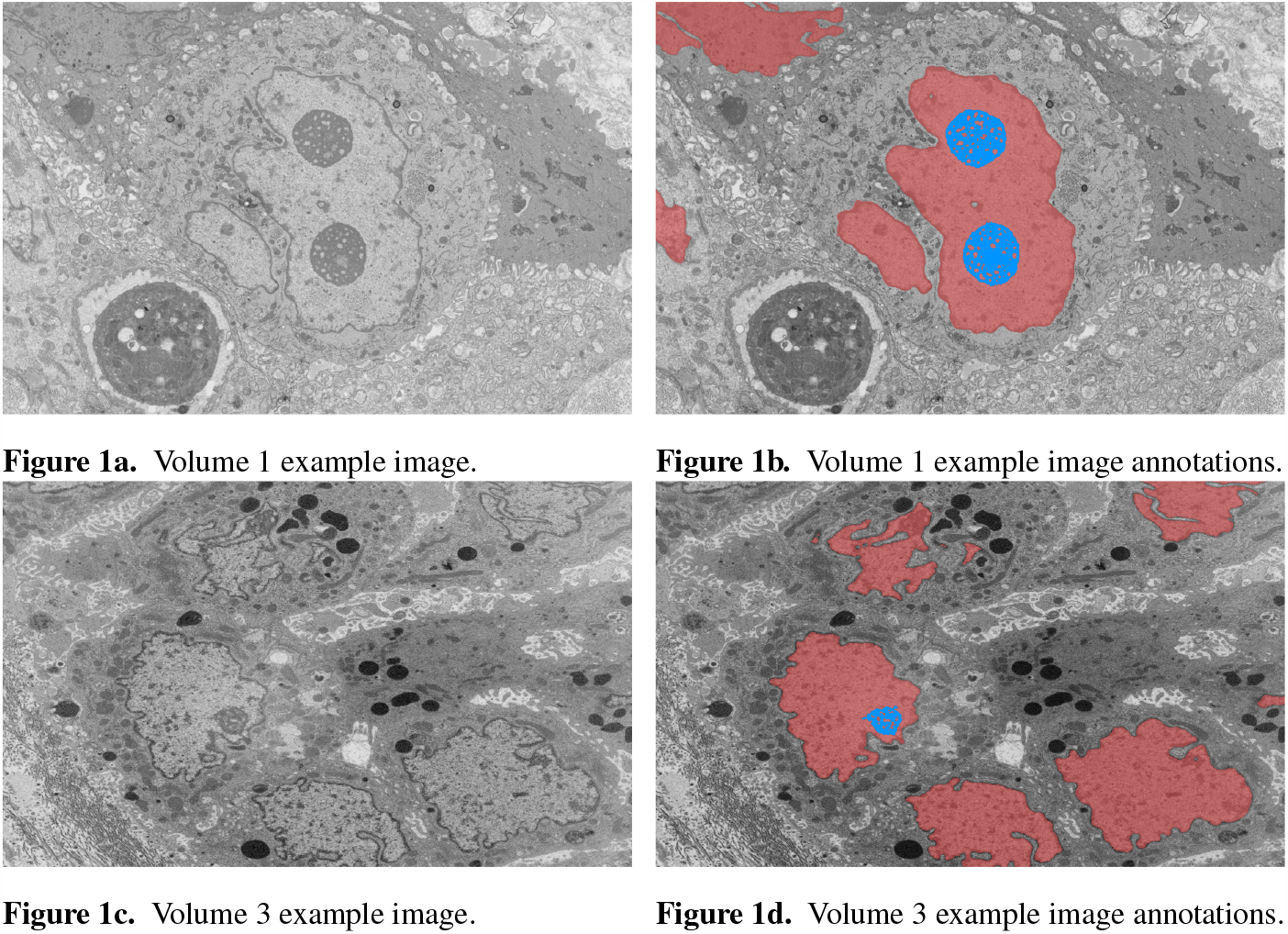
Example image slices and corresponding ground truth annotations from Volumes 1 and 3. The horizontal image width is equal to 25 μm. Nuclei are in red and nucleoli in blue. **(A)** Volume 1 example image. **(B)** Volume 1 example image annotations. **(C)** Volume 3 example image. **(D)** Volume 3 example image annotations.

**Figure 2.**
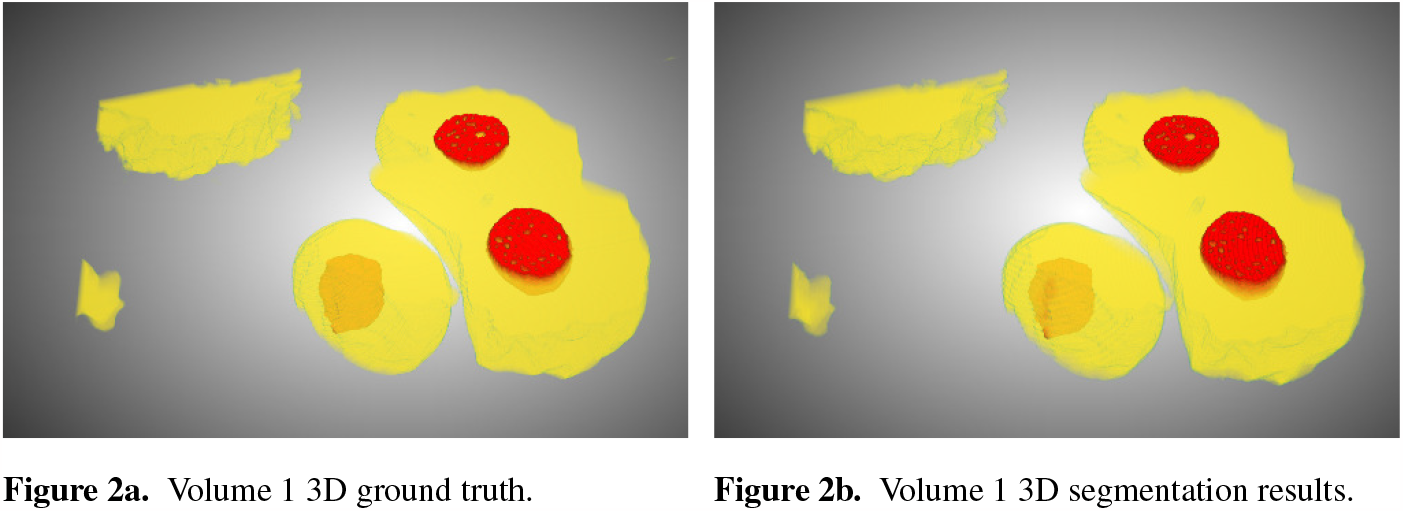
Ground truth (a) and segmentation by SSL-UNet++-CutMix (b) 3D visualizations for Volume 1. Nuclei are in yellow and nucleoli in red.

## 2 MATERIALS AND METHODS

### 2.1 Training and evaluation

Previously, we trained a model for each dataset using a subset of manually labeled images spaced evenly along the volumes, and evaluated on remaining unlabeled images. We reported results on using 7, 10, 15 and 25 training images on all datasets, which represents between 0.3% and 3.3% of all image slices depending on the dataset. As these EM images had large dimensions (typically around 6000 *×* 4000 pixels), they were cropped to 512 *×* 512 tiles. We followed the same procedure as (Machireddy et al., 2021) of extracting tiles of size 2048 *×* 2048 and down sampling them to 512 *×* 512 as a way to artificially add context. We applied standard random flip and rotation data augmentations. When training with nucleoli, as they account for a small area in the total image, taking random crops effectively resulted in most crops being empty, and models collapsing to the prediction of background. To address this issue, we ensured that more than 99% of crops in a batch contain nucleoli.

Moreover, we selected the Dice score as an evaluation metric because it can be seen as a harmonic mean of precision and recall, and is fitted when dealing with imbalanced classes setting, which is our case. However, we also report the 3D Dice score rather than the average of individual Dice scores across all slices as reported in (Machireddy et al., 2021). Indeed, we found the latter to be biased towards giving more importance to slices with fewer foreground pixels, while the former effectively reflects the captured percentage of the target structure. For the sake of comparison we reported the averaged version in our results section in addition to the 3D Dice scores. We recommend however to use the latter. The 3D Dice and average dice are more precisely defined as follows:

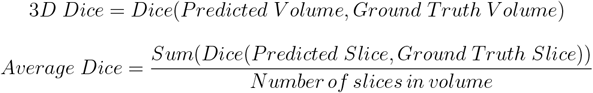

All models were trained and evaluated on one NVIDIA V100 GPU, as we strongly believe we should keep our clinical end-goal in mind and aim to reflect image analysis capabilities available to teams with reasonable computational power. To this end, we also report training times and number of parameters in table 1.

**Table 1.** Number of parameters and training times for one volume. SSL trained models require double the number of parameters because two models are trained at the same time.

|  | UNet++ | FracTALResNet | CEECNet | Senformer | SSL-ResUNet | SSL-UNet++ |
| --- | --- | --- | --- | --- | --- | --- |
| # of parameters | 26,072,337 | 18,199,919 | 58,964,079 | 163,100,906 | 4,283,201 * 2 | 21,954,705 * 2 |
| Average training time | ~48 hours | ~70 hours | ~90 hours | ~144 hours | ~60 hours | ~64 hours |

### 2.2 Fully-supervised framework

#### 2.2.0.1 ResUNet

We used previous work from (Machireddy et al., 2021) as the baseline for nuclei and nucleoli segmentation. The model used was a Residual U-Net (ResUNet) (He et al., 2015), a simple yet robust fully convolutional encoder-decoder network. U-net and its variants are the most prominent architectures for image segmentation, as the residual connections solve the gradient vanishing problem faced when working with very deep models (Glorot and Bengio, 2010), while the different levels allow feature refinement at different scales. These features have made U-nets widely used in many computer vision problems, including analysis of medical data (Su et al., 2021; Siddique et al., 2020).

#### 2.2.0.2 UNet++

UNet++ (Zhou et al., 2018) was the first model we decided to compare to the baseline. Our motivation to use this model came from the fact that it was heavily inspired by ResUNet, and was especially designed for medical-like image segmentation. The major differences from ResUNet are the presence of dense convolution blocks on the skip connections and deep supervision losses. Dense convolution blocks aim at reducing the semantic gap between the encoder and decoder, while deep supervision loss enables the model to be accurate (by averaging outputs from all segmentation branches) and fast (by selecting one of the segmentation maps as output). In our work, as we were primarily focused on accuracy, we used the average of all branches. We used the implementation available in the *Segmentation Models* Python library ^1^ with ResNet34 as the encoder backbone, and the soft Dice loss (DL) function which is commonly used in semantic segmentation for images when background and foreground classes are imbalanced, and is defined as follows for a ground truth *y* and prediction 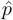:

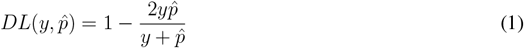

#### 2.2.0.3 FracTALResNet

FracTALResNet (Diakogiannis et al., 2021) was also used for comparison. While the original model presented is designed for the task of semantic change detection, it can be adapted for semantic segmentation, and such architecture is in fact available in the authors’ official implementation ^2^. It was heavily inspired from ResUNet as well, but makes use of a multi-head attention layer (FracTAL block). It also makes use of boundaries and distance maps calculated from the segmentation masks in order to improve performances, but at the cost of both memory and computational time during training. It is trained using the Fractal Tanimoto similarity measure.

#### 2.2.0.4 CEECNet

CEECNet was also introduced in (Diakogiannis et al., 2021) and for the same purpose as FracTALResNet, but managed to achieve state-of-the-art performances by focusing on context. Indeed, the CEECNet block stands for Compress-Expand Expand-Compress and is comprised of two branches. The first branch (CE block) processes a view of the input in lower resolution, while the second branch (EC block) treats a view in higher spatial resolution. Motivation behind using this model came from the fact that, as described in section 2.1, feeding more context by down-sampling to a lower resolution is beneficial to segmentation accuracy. Since the core block of CEECNet is based on the compress and expand operations, we believed this network would be able to leverage contextual information in order to achieve better segmentation performances. Similar to FracTALResNet, it was trained with the Tanimoto similarity measure and needs computed boundaries and distance maps.

#### 2.2.0.5 SenFormer

SenFormer (Bousselham et al., 2021) (Efficient Self-Ensemble Framework for Semantic Segmentation) was the last fully-supervised method tested. It is a newly developed ensemble approach for the task of semantic segmentation that makes use of transformers in the decoders and the Feature Pyramid Network (FPN) backbone. Our motivation behind using this model came from the fact that it is almost purely attention-based, which by definition adds spatial context to the segmentation.

#### 2.2.0.6 Supervised model choice

Since model architecture is orthogonal to using a semi-supervised framework, we picked the best performing model using Dice scores, as can be seen in Table 2. UNet++ performed better on average, exhibited a low variance, often performed the best out of fully-supervised architectures, and almost never under performed (as shown by the average rank). Detailed results were reported in Tables 3 and 4. For these reasons, it was the model we chose to compare to the baseline in the semi-supervised framework.

**Table 2.** Average, standard deviation and average rank for Dice score over all volumes.

|  | ResUNet | UNet++ | FracTALResNet | CEECNet | Senformer |
| --- | --- | --- | --- | --- | --- |
| Average | 0.9200 | **0.9270** | 0.9250 | 0.9036 | 0.9146 |
| Standard deviation | 0.0107 | 0.0083 | **0.0072** | 0.137 | 0.0099 |
| Average rank | 2.625 | **2.6042** | 3.1875 | 3.54 | 3.0417 |

**Table 3.** Nuclei segmentation Dice scores. Columns labeled 7, 10, 15 and 25 in the second line represent the number of training images for each volume.

|  | Volume 1 |  |  |  | Volume 2 |  |  |  | Volume 3 |  |  |  |
| --- | --- | --- | --- | --- | --- | --- | --- | --- | --- | --- | --- | --- |
|  | 7 | 10 | 15 | 25 | 7 | 10 | 15 | 25 | 7 | 10 | 15 | 25 |
| ResUNet | 0.9805 | 0.9846 | 0.9846 | 0.9847 | 0.9845 | 0.9875 | 0.9878 | 0.9933 | 0.9597 | 0.9737 | 0.9738 | 0.9745 |
| UNet++ | 0.9746 | 0.9791 | 0.9801 | 0.9844 | 0.988 | 0.9908 | 0.9922 | 0.993 | 0.9606 | 0.9625 | 0.9705 | 0.985 |
| FracTALResNet | 0.9724 | 0.9796 | 0.9817 | 0.9885 | 0.9756 | 0.9836 | 0.9871 | 0.9887 | 0.9655 | 0.9698 | 0.976 | 0.9825 |
| CEECNet | 0.9702 | 0.9688 | 0.9825 | 0.9742 | 0.9847 | 0.9894 | 0.9928 | 0.9948 | 0.9457 | 0.9499 | 0.9509 | 0.98 |
| Senformer | 0.9821 | 0.9851 | 0.9877 | 0.9897 | 0.9835 | 0.987 | 0.9898 | 0.9927 | 0.971 | 0.9722 | 0.979 | 0.9858 |
| SSL-ResUNet | 0.988 | 0.9889 | 0.9882 | 0.9902 | 0.9938 | 0.9931 | 0.9941 | 0.9958 | 0.9703 | 0.9702 | 0.977 | 0.9822 |
| SSL-ResUNet-CutMix | 0.9892 | 0.9898 | 0.9903 | 0.9912 | **0.9951** | **0.9952** | **0.9954** | 0.9957 | 0.9726 | 0.9747 | 0.9799 | 0.9822 |
| SSL-UNet++-CutMix | **0.9903** | **0.9910** | **0.9912** | **0.9923** | **0.9951** | **0.9952** | **0.9954** | **0.9959** | **0.9804** | **0.9839** | **0.9859** | **0.9872** |

**Table 4.** Nucleoli segmentation Dice scores. Columns labeled 7, 10, 15 and 25 in the second line represent the number of training images for each volume.

|  | Volume 1 |  |  |  | Volume 2 |  |  |  | Volume 3 |  |  |  |
| --- | --- | --- | --- | --- | --- | --- | --- | --- | --- | --- | --- | --- |
|  | 7 | 10 | 15 | 25 | 7 | 10 | 15 | 25 | 7 | 10 | 15 | 25 |
| ResUNet | 0.9686 | 0.9733 | 0.9664 | 0.9712 | 0.8811 | 0.9019 | 0.9108 | 0.9182 | 0.7054 | 0.7014 | 0.6906 | 0.7166 |
| UNet++ | 0.9576 | 0.9624 | 0.9652 | 0.9671 | 0.8957 | 0.9168 | 0.9139 | 0.9216 | 0.638 | 0.7321 | 0.7893 | 0.8282 |
| FracTALResNet | 0.9662 | 0.9613 | 0.9464 | 0.9671 | 0.8809 | 0.8778 | 0.8775 | 0.9237 | 0.6892 | 0.7547 | 0.7677 | 0.8307 |
| CEECNet | 0.9779 | 0.9775 | 0.9772 | **0.9797** | 0.7615 | 0.8608 | 0.8339 | 0.8565 | 0.586 | 0.6567 | 0.7321 | 0.8036 |
| Senformer | 0.9353 | 0.9384 | 0.9413 | 0.9442 | 0.8385 | 0.8594 | 0.8818 | 0.907 | 0.6695 | 0.6678 | 0.7538 | 0.8084 |
| SSL-ResUNet | 0.9775 | **0.9783** | 0.9767 | 0.9782 | 0.9007 | 0.9196 | 0.9344 | 0.9401 | 0.6714 | 0.7447 | 0.8041 | 0.8176 |
| SSL-ResUNet-CutMix | **0.9781** | 0.9782 | **0.9787** | 0.9791 | 0.9193 | 0.9218 | 0.9339 | 0.9440 | 0.7763 | 0.8013 | 0.8159 | 0.8266 |
| SSL-UNet++-CutMix | 0.9777 | **0.9783** | 0.9781 | 0.9793 | **0.9292** | **0.9256** | **0.9349** | **0.9473** | **0.7887** | **0.8071** | **0.8240** | **0.8390** |

### 2.3 Semi-supervised learning (SSL) framework

#### 2.3.0.1 Cross Pseudo Supervision (CPS)

As described in section 2.1, roughly 1% of the collected images were manually annotated and used for training the fully-supervised methods. To take advantage of the potential semantic information contained in the unlabeled images, we used the CPS framework described in (Chen et al., 2021). It trained two networks with a standard supervised cross-entropy loss and used pseudo-labels generated from the segmentation confidence map of one network to supervise the other as can be seen in Figure 3. Loss for unlabeled images in CPS is defined as follows with *D*^*u*^ denoting unlabeled data, *p*_*i*_ the segmentation confidence map, *y*_*i*_ the predicted label map, *ℓ*_*ce*_ the cross-entropy loss, 1, and 2 representing each network:

**Figure 3.**
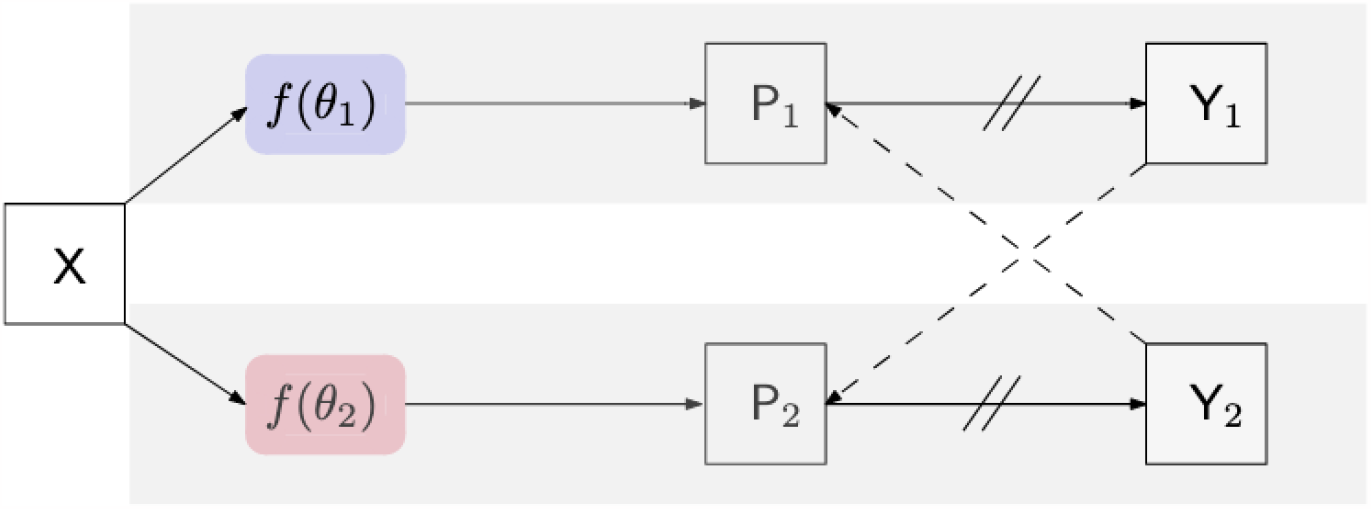
Illustration of the CPS training framework. *P*_*i*_ is the segmentation confidence map, *Y*_*i*_ is the predicted label map. *→* means forward operation, --→ loss supervision, and // on *→* stopping the gradient.

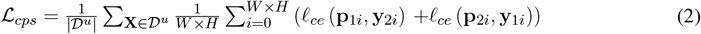

We trained both ResUNet and UNet++ models with this framework. In this benchmark, we trained models with the soft Dice loss defined in equation 1, as we noticed that models trained with a loss closer to the evaluation metric performed better. As a consequence, we replaced all of the cross-entropy losses originally used in (Chen et al., 2021) by the soft Dice loss. We also noticed that learning needed to be driven by the supervised loss during the first epochs. At the beginning of training, models had no prior knowledge of the segmentation task, and thus, could not yield relevant pseudo-labels resulting in frequent collapsing to predict only the background, especially when working with nucleoli. To resolve this, we implemented a linear warm-up to *λ*, the parameter used to balance the CPS loss with the supervised loss, so that the latter has priority over the former during early steps of training. We used a value of 1 for *λ* in all of our experiments.

#### 2.3.0.2 Integration of CutMix data augmentation

CutMix (Yun et al., 2019) is a popular data augmentation method for training classifiers that shuffles information throughout the training batch, and has recently been used in semi-supervised segmentation tasks. In the authors’ implementation, when using the CutMix strategy in the CPS loss, the latter is only optionally computed on labeled data. However, in our case, not having labels meant not being able to ensure the CPS batch contained any nucleoli as we did for supervised methods (see section 2.1). This made the loss unstable as models performed poorly on empty images. To solve this issue, we trained all models with both the supervised and unsupervised CPS loss, and ensured that at least half of the CPS batch contained nucleoli. We tried using CutMix in the fully supervised setting, however, it did not yield any significant improvement. We believe this to be due to the fact that the number of images we trained on was so limited in the fully-supervised setting, CutMix could not add much new information during augmentation.

## 3 RESULTS AND DISCUSSION

### 3.0.0.1 Accurate segmentation of nuclei and nucleoli

For comparison with previous results (Machireddy et al., 2021), we first used the same average of 2D dices (tables 3,4) while also providing the unbiased 3D Dice metric results (tables 5,6). 3D dice scores are almost always higher than their average counterpart, which is positive as they are more representative of the segmentation quality. This is particularly visible for nucleoli (compare tables 4 and 6). As can be observed in Tables 3 and 4, all evaluated models were able to accurately segment both nuclei and nucleoli. Despite the introduction of attention enabling models to gain a marginal edge over the baseline introduced in (Machireddy et al., 2021), a clear advantage was obtained only when using SSL, especially when paired with CutMix data augmentation. The best results are given by UNet++ with SSL and CutMix, which indicates both model selection and using a semi-supervised framework help improve segmentation performance.

**Table 5.**
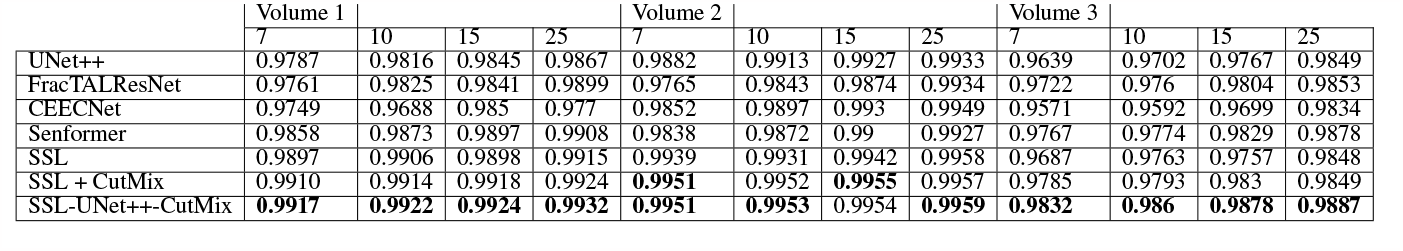
3D nuclei segmentation Dice scores. Numbers 7, 10, 15 and 25 in the second line represent the number of training images for each volume.

**Table 6.** 3D nucleoli segmentation Dice scores. Numbers 7, 10, 15 and 25 in the second line represent the number of training images for each volume.

|  | Volume 1 |  |  |  | Volume 2 |  |  |  | Volume 3 |  |  |  |
| --- | --- | --- | --- | --- | --- | --- | --- | --- | --- | --- | --- | --- |
|  | 7 | 10 | 15 | 25 | 7 | 10 | 15 | 25 | 7 | 10 | 15 | 25 |
| UNet++ | 0.9626 | 0.9647 | 0.9664 | 0.9681 | 0.9073 | 0.9321 | 0.9333 | 0.9362 | 0.734 | 0.8115 | 0.8438 | 0.869 |
| FracTALResNet | 0.9755 | 0.9657 | 0.9522 | 0.9685 | 0.8814 | 0.8722 | 0.8761 | 0.9371 | 0.7523 | 0.8044 | 0.8365 | 0.8672 |
| CEECNet | 0.9788 | 0.979 | 0.979 | **0.9806** | 0.7852 | 0.8902 | 0.8436 | 0.8899 | 0.6629 | 0.7186 | 0.7927 | 0.8419 |
| Senformer | 0.9396 | 0.9398 | 0.9425 | 0.9453 | 0.8585 | 0.8734 | 0.8962 | 0.9144 | 0.7556 | 0.7778 | 0.8231 | 0.8473 |
| SSL | 0.9789 | 0.9797 | 0.9782 | 0.9792 | 0.9109 | 0.9152 | 0.9436 | 0.9513 | 0.7662 | 0.806 | 0.8545 | 0.8614 |
| SSL + CutMix | **0.9797** | 0.9797 | **0.98** | 0.9801 | 0.9167 | **0.9231** | 0.946 | 0.9525 | 0.8235 | 0.8415 | 0.8558 | 0.8667 |
| SSL-UNet++-CutMix | 0.9794 | **0.9798** | 0.9793 | 0.9803 | **0.9415** | 0.9198 | **0.9475** | **0.9536** | **0.8421** | **0.8436** | **0.8626** | **0.8726** |

### 3.0.0.2 Benefits of semi-supervised learning

While fully-supervised models could sometimes outperform SSL ones on specific datasets (for example CEECNet on Volume 1 nucleoli), SSL remained stable over all structures and volumes. It outperformed the baseline for all datasets, most noticeably on Volume 3 nucleoli, with an average gain of 0.11 in Dice, representing a 15.6% performance increase. One of the reasons behind this performance gain is the high heterogeneity in Volume 3 nucleoli, and most models struggled segmenting unseen structures, as can be observed in Figure 4. As the performances of different fully-supervised methods varied highly depending on the volumes (for example Senformer under-performed in segmenting Volume 1 nucleoli), the SSL methods remained stable.

**Figure 4.**
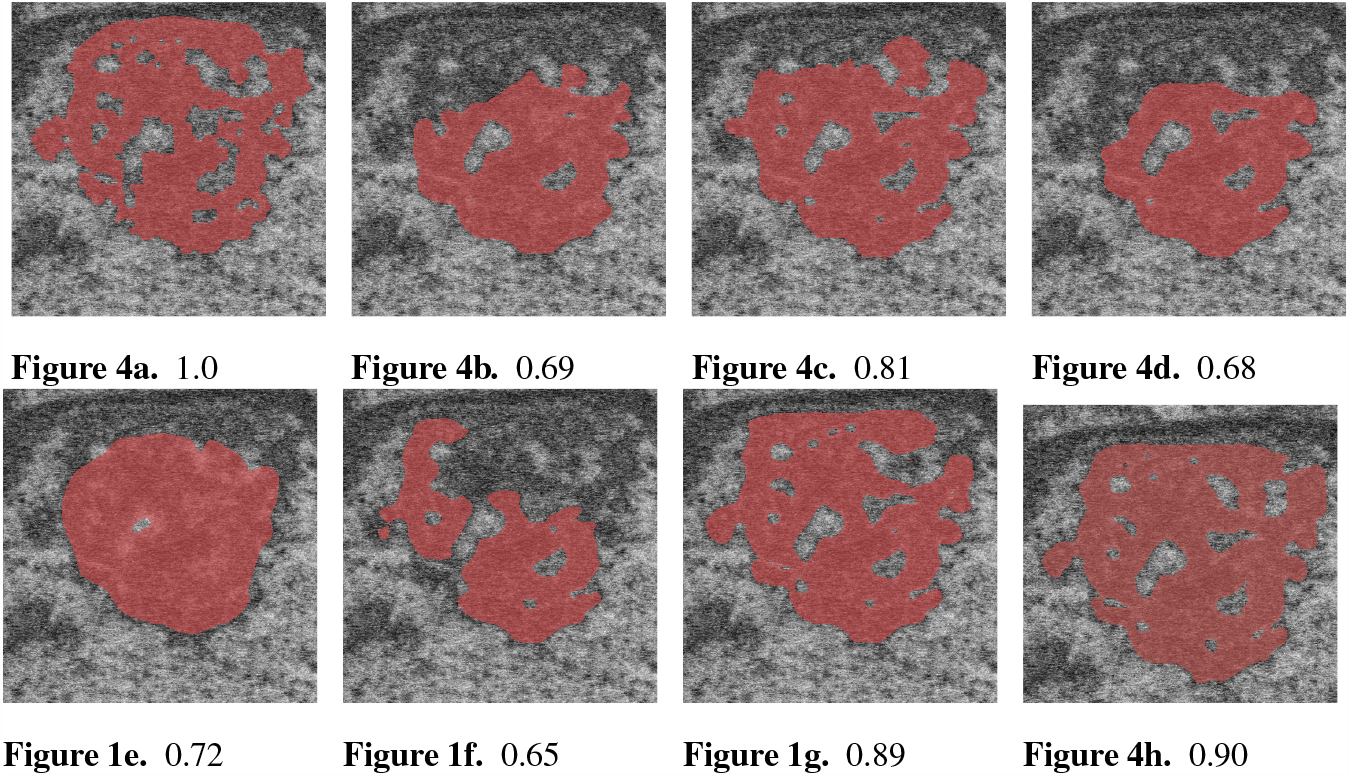
Qualitative results with Dice score for a difficult nucleolus in Volume 3, from (a) ground truth, (b) UNet++, (c) FracTALResNet, (d) CEECNet, (e) Senformer, (f) SSL-ResUNet, (g) SSL-ResUNet-CutMix and (h) SSL-UNet++-CutMix

When evaluating our models, we noticed that fully supervised methods performed really well around the images they were trained on (see Figure 5), yielding a near perfect Dice score. However, performances dropped as soon as evaluation images start being dissimilar to the training images, thus forming dips visible in the plot. This is a clear sign of over-fitting that SSL prevented thanks to the regularization added by the CPS loss. This stability and consistency across image volumes allows, in addition to the performance gain, an easier post-processing of the segmented volume by manual inspection and interpretation or algorithmic analysis. These result made us believe semi-supervised frameworks were key in attaining better generalization performance in our sparse annotation setting. Indeed, the Dice score of UNet++ with SSL and Cutmix are better most of the time with only 7 training images than what was achieved previously with 25 images in (Machireddy et al., 2021) or with supervised models in this study.

**Figure 5.**
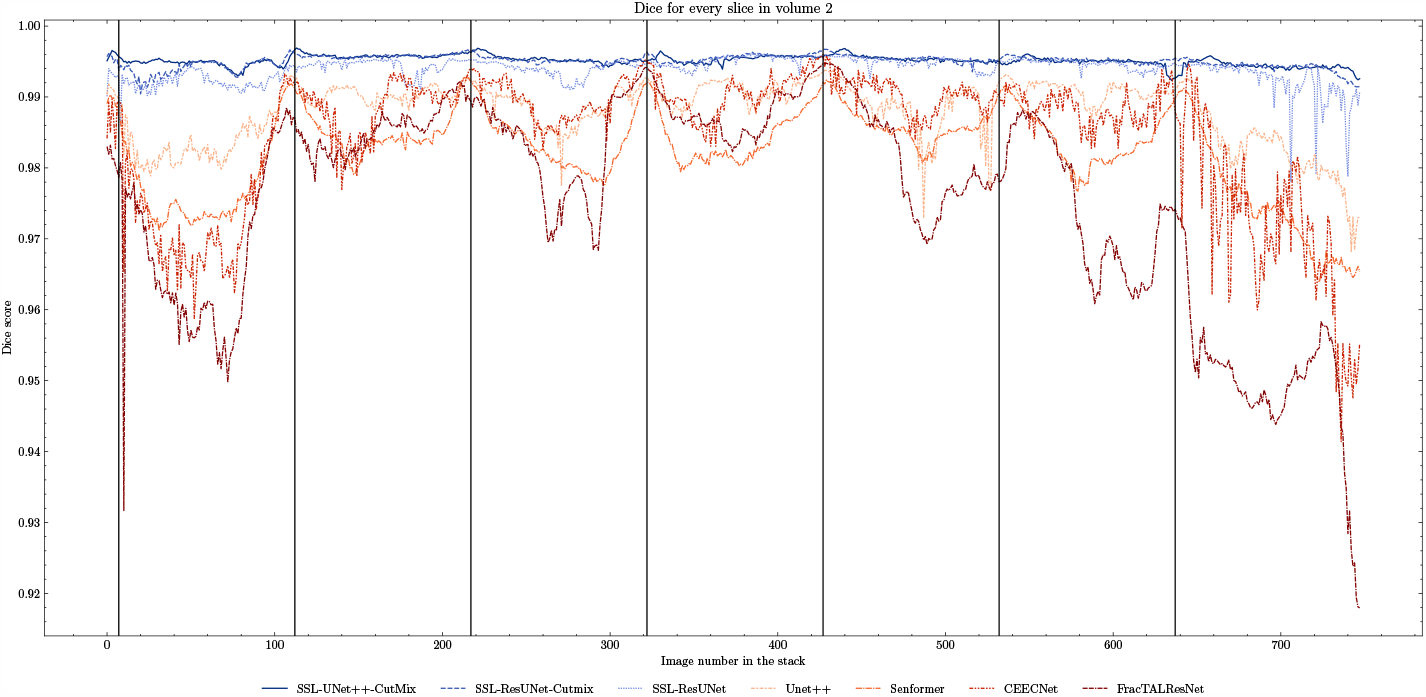
Comparison of Dice scores for nuclei segmentation along all 757 slices in Volume 2 with 7 training images. Training slices are marked with vertical black lines. We can clearly observe the 7 peaks in performance and drops in-between for fully-supervised methods (beige to brown) as opposed to the stability of the SSL models (blue).

## 4 CONCLUSION

In this work, we investigated the segmentation of nuclei and nucleoli in vEM images of cancer cells. We studied the performances of several leading deep learning models and assessed the relative performance gains of each method. We provided insight as to why semi-supervised methods were able to yield more robust results and managed to improve on previous work both in terms of reducing the amount of data needed and segmentation performances, with an improved Dice on all Volumes. We made the experiment code available at ^3^ and the complete manual annotations for the data have been provided through the HTAN data portal. We believe that semi-supervised methods are a key component in segmentation with sparse annotations as they proved to be superior in both quantitative and qualitative evaluations.

## AUTHOR CONTRIBUTIONS

Conceptualization, J.W.G; Methodology, L.P, G.T and X.S; Software, L.P, W.B and G.T; Formal Analysis, L.P, X.S, G.T; Investigation, L.P, X.S, J.L.R. and G.T; Data Curation, J.L.R; Writing – Original Draft, L.P and X.S; Writing – Review & Editing, L.P, X.S, G.T, J.L.R and W.B; Visualization, L.P; Supervision, X.S, G.T, J.L.R and J.W.G.; Project Administration, J.W.G; Funding Acquisition, J.W.G.

## ACKNOWLEDGMENTS

FIB-SEM data included in this manuscript were generated at the Multiscale Microscopy Core (MMC), an OHSU University Shared Resource, with technical support from the OHSU Center for Spatial Systems Biomedicine (OCSSB). The authors acknowledge the Knight Cancer Institute’s Precision Oncology SMMART Clinical Trials Program for the resources, samples, and data that supported this study. Specimen acquisition support from the SMMART clinical coordination team was invaluable. This work is supported by the Cancer Early Detection Advanced Research (CEDAR) Center at the Knight Cancer Institute at OHSU. This study was approved by the Oregon Health and Science University Institutional Review Board (IRB16113). Participant eligibility was determined by the enrolling physician and informed consent was obtained prior to all study protocol related procedures.

## DATA AVAILABILITY STATEMENT

The datasets analyzed for this study and corresponding ground truths will be provided upon request. The three datasets from breast carcinoma tissue are available through the HTAN Data Coordinating Center (https://humantumoratlas.org/) as patient HTA9-1 in the OMS Atlas. HTAN sample IDs for Bx1, Bx2 and Bx4 are HTA9-1 bx1, HTA9-1 bx2 and HTA9-5 (bx4), respectively.

## Footnotes

1 https://github.com/qubvel/segmentation_models.pytorch

2 https://github.com/feevos/ceecnet

3 https://github.com/LucasPagano/Segmentation3DEM

## REFERENCES

Baghban, R., Roshangar, L., Jahanban-Esfahlan, R., Seidi, K., Ebrahimi-Kalan, A., Jaymand, M., et al. (2020). Tumor microenvironment complexity and therapeutic implications at a glance. Cell Commun Signal 18, 59

Bousselham, W., Thibault, G., Pagano, L., Machireddy, A., Gray, J., Chang, Y. H., et al. (2021). Efficient self-ensemble framework for semantic segmentation

Bushby, A. J. P K. M. Y., Young, R. D., Pinali, C., Knupp, C., and Quantock, A. J. (2011). Imaging three-dimensional tissue architectures by focused ion beam scanning electron microscopy doi:10.1038/nprot.2011.332

Chen, X., Yuan, Y., Zeng, G., and Wang, J. (2021). Semi-supervised semantic segmentation with cross pseudo supervision, 2613–2622doi:10.1109/cvpr46437.2021.00264

Diakogiannis, F. I., Waldner, F., and Caccetta, P. (2021). Looking for change? roll the dice and demand attention. Remote Sensing 2021, Vol. 13, Page 3707 13, 3707. doi:10.3390/RS13183707

Giannuzzi, L. A. and Stevie, F. A. (2005). Introduction to focused ion beams

Glorot, X. and Bengio, Y. (2010). Understanding the difficulty of training deep feedforward neural networks. Journal of Machine Learning Research - Proceedings Track 9, 249–256

[Dataset] He, K., Zhang, X., Ren, S., and Sun, J. (2015). Deep residual learning for image recognition

Hirata, E. and Sahai, E. (2017). Tumor Microenvironment and Differential Responses to Therapy. Cold Spring Harb Perspect Med 7

Jain, J., Singh, A., Orlov, N., Huang, Z., Li, J., Walton, S., et al. (2021). Semask: Semantically masked transformers for semantic segmentation

Karabağ, C., Jones, M. L., Peddie, C. J., Weston, A. E., Collinson, L. M., and Reyes-Aldasoro, C. C. (2020). Semantic segmentation of hela cells: An objective comparison between one traditional algorithm and four deep-learning architectures. PLoS ONE 15. doi:10.1371/JOURNAL.PONE.0230605

Lindström, M. S., Jurada, D., Bursac, S., Orsolic, I., Bartek, J., and Volarevic, S. (2018). Nucleolus as an emerging hub in maintenance of genome stability and cancer pathogenesis. Oncogene 37, 2351–2366. doi:10.1038/s41388-017-0121-z

Machireddy, A., Thibault, G., Loftis, K. G., Stoltz, K., Bueno, C. E., Smith, H. R., et al. (2021). Robust segmentation of cellular ultrastructure on sparsely labeled 3d electron microscopy images using deep learning. bioRxiv doi:10.1101/2021.05.27.446019

Nunney, L., Maley, C. C., Breen, M., Hochberg, M. E., and Schiffman, J. D. (2015). Peto’s paradox and the promise of comparative oncology. Philosophical Transactions of the Royal Society B: Biological Sciences 370. doi:10.1098/rstb.2014.0177

Perez, A. J., Seyedhosseini, M., Deerinck, T. J., Bushong, E. A., Panda, S., Tasdizen, T., et al. (2014). A workflow for the automatic segmentation of organelles in electron microscopy image stacks. Frontiers in Neuroanatomy 8. doi:10.3389/FNANA.2014.00126/ABSTRACT

Shi, J., Kantoff, P. W., Wooster, R., and Farokhzad, O. C. (2017). Cancer nanomedicine: progress, challenges and opportunities. Nat Rev Cancer 17, 20–37

Siddique, N., Paheding, S., Elkin, C. P., and Devabhaktuni, V. (2020). U-net and its variants for medical image segmentation: theory and applications. IEEE Access doi:10.1109/access.2021.3086020

Su, R., Zhang, D., Liu, J., and Cheng, C. (2021). Msu-net: Multi-scale u-net for 2d medical image segmentation. Frontiers in Genetics 12, 140. doi:10.3389/FGENE.2021.639930/BIBTEX

Taha, A. A. and Hanbury, A. (2015). Metrics for evaluating 3d medical image segmentation: analysis, selection, and tool. BMC Medical Imaging 15, 29. doi:10.1186/s12880-015-0068-x

Tanaka, H. Y. and Kano, M. R. (2018). Stromal barriers to nanomedicine penetration in the pancreatic tumor microenvironment. Cancer Sci 109, 2085–2092

Yun, S., Han, D., Chun, S., Oh, S. J., Choe, J., and Yoo, Y. (2019). Cutmix: Regularization strategy to train strong classifiers with localizable features. Proceedings of the IEEE International Conference on Computer Vision 2019-October, 6022–6031. doi:10.1109/ICCV.2019.00612

Zhang, F., Breger, A., Cho, K. I. K., Ning, L., Westin, C. F., O’Donnell, L. J., et al. (2021). Deep learning based segmentation of brain tissue from diffusion mri. NeuroImage 233, 117934. doi:10.1016/J.NEUROIMAGE.2021.117934

Zhou, Z., Siddiquee, M. M. R., Tajbakhsh, N., and Liang, J. (2018). Unet++: A nested u-net architecture for medical image segmentation. CoRR abs/1807.10165

Zink, D., Fischer, A. H., and Nickerson, J. A. (2004). Nuclear structure in cancer cells. Nature reviews.Cancer 4, 677–687. doi:10.1038/NRC1430

